# Predicting motivation: computational models of PFC can explain neural coding of motivation and effort-based decision-making in health and disease

**DOI:** 10.1101/171637

**Authors:** Eliana Vassena, James Deraeve, William H. Alexander

## Abstract

Human behavior is strongly driven by the pursuit of rewards. In daily life, however, benefits mostly come at a cost, often requiring that effort be exerted in order to obtain potential benefits. Medial prefrontal cortex (MPFC) and dorsolateral prefrontal cortex (DLPFC) are frequently implicated in the expectation of effortful control, showing increased activity as a function of predicted task difficulty. Such activity partially overlaps with expectation of reward, and has been observed both during decision-making and during task preparation. Recently, novel computational frameworks have been developed to explain activity in these regions during cognitive control, based on the principle of prediction and prediction error (PRO model, Alexander and Brown, 2011, HER Model, Alexander and Brown, 2015). Despite the broad explanatory power of these models, it is not clear whether they can also accommodate effects related to the expectation of effort observed in MPFC and DLPFC. Here, we propose a translation of these computational frameworks to the domain of effort-based behavior. First, we discuss how the PRO model, based on prediction error, can explain effort-related activity in MPFC, by reframing effort-based behavior in a predictive context. We propose that MPFC activity reflects monitoring of motivationally relevant variables (such as effort and reward), by coding expectations, and discrepancies from such expectations. Moreover, we derive behavioral and neural model-based predictions for healthy controls and clinical populations with impairments of motivation. Second, we illustrate the possible translation to effort-based behavior of the HER model, an extended version of PRO model based on hierarchical error prediction, developed to explain MPFC-DLPFC interactions. We derive behavioral predictions which describe how effort and reward information is coded in PFC, and how changing the configuration of such environmental information might affect decision-making and task-performance involving motivation.

## Introduction

Attaining a goal often requires commitment, implementation of a precise course of actions, and deployment of sufficient resources to reach successful completion. How humans fulfill this process is largely investigated in the field of cognitive neuroscience under the term of goal-directed behavior. A crucial underlying cognitive mechanism is prediction: the ability to evaluate the environment, formulate expectations about future events based on previous experiences, and finally compare such expectations with subsequent outcomes in order to update one’s knowledge about the current state of the world. Furthermore, adaptive interaction with the environment entails predicting the impact of one’s action and evaluating outcomes.

The human Prefrontal Cortex (PFC) is the neural machinery that supports crucial mechanisms involved in goal-directed behavior (Miller & Cohen, 2001). In particular, the medial portion of PFC (MPFC, including dorsal Anterior Cingulate Cortex, dACC) is implicated in prediction and outcome evaluation (Alexander & Brown, 2011; Jahn, Nee, Alexander, & Brown, 2014), including situations in which the outcome carries value for the agent, such as a reward (Rushworth, Walton, Kennerley, & Bannerman, 2004; Silvetti, Seurinck, & Verguts, 2011, 2013; Vassena, Krebs, Silvetti, Fias, & Verguts, 2014). Notably, a variety of other functions have been attributed to MPFC (Vassena, Holroyd, & Alexander, 2017), including error monitoring, (Holroyd, Nieuwenhuis, Mars, & Coles, 2004; van Veen, Holroyd, Cohen, Stenger, & Carter, 2004), conflict detection (Botvinick, Braver, Barch, Carter, & Cohen, 2001), pain and affect processing (Bush, Luu, & Posner, 2000; Nee, Kastner, & Brown, 2011), and value-based decision-making (Rangel & Hare, 2010; Rushworth, Kolling, Sallet, & Mars, 2012; Rushworth & Behrens, 2008). Recent computational work has explained this array of empirical findings under the unifying principle of prediction and prediction error. In this framework, MPFC formulates predictions tracking stimuli, actions and outcomes, and computes a signal termed prediction error, which scales with the discrepancy between predicted and actual outcomes (Predicted Response Outcome model, PRO model, Alexander & Brown, 2011). This mechanism allows rapid prediction updating according to environmental feedback, be it an error, a painful stimulus, or a reward. An extended version of the same model, the Hierarchical Error Representation model (HER, Alexander & Brown 2015), expands the same computational principle in a hierarchical architecture, capturing more complex high-level cognitive processes involving the interaction of MPFC and dorsolateral PFC (DLPFC), typically associated with higher-level cognitive functions such as working memory and goal-maintenance (Miller & Cohen, 2001).

The goal of this manuscript is to explore the power of these computational accounts, in terms of generating novel neural and behavioral predictions for untested contexts and populations. These frameworks have proven useful across several fields of cognition, yet they have not been put to test in the field of effortful behavior and motivation. Goal-directed behavior generally involves competing factors, including the value of the prospective goals, how much effort one is willing to exert to attain the desired goal, and preparation for the necessary effortful performance (Botvinick & Braver, 2015; Westbrook & Braver, 2013, 2015). First, we will describe MPFC involvement in effort-based behavior. Then, we illustrate how the PRO model can be generalized to the domain of motivation. We propose that MPFC activity reflects monitoring of motivationally relevant variables such as reward and required effort, instead of coding an explicit cost-benefit or choice signal per se. We illustrate novel model-based simulations, as well as theoretical predictions, which can be used to guide further empirical enquiry. We discuss how the PRO framework makes neural and behavioral predictions for clinical conditions in which motivation is impaired, such as depression and other psychiatric disorders (Treadway, Bossaller, Shelton, & Zald, 2012). Subsequently, we discuss the future directions in translating the HER model to the domain of motivation, extrapolating behavioral predictions.

From such predictions, we derive implications for measuring and potentially training motivation-related cognitive mechanisms in clinical populations.

## Effort-based decision-making and performance in MPFC

Experimental manipulation of effort in behavioral and neuroimaging experiments has yielded a wealth of findings in the past decade. Typically, effort is perceived as aversive (Kool, McGuire, Rosen, & Botvinick, 2010), yet humans decide to engage in it when doing so leads to a benefit, such as a reward. In the framework of decision-making and neuroeconomics, the net value of a potential reward is discounted (i.e. decreased) by the amount of effort required to obtain the reward (Apps, Grima, Manohar, & Husain, 2015; Hartmann, Hager, Tobler, & Kaiser, 2013; Nishiyama, 2014). This seems to hold across different types of effort (Nishiyama, 2016), and guides decisions to engage one’s resources in the task at hand (Kurzban, Duckworth, Kable, & Myers, 2013). Several studies in animals described the neural mechanisms underlying this cost-benefit evaluation, showing a pivotal role of MPFC in interaction with striatal and sub-cortical nuclei (Hosokawa, Kennerley, Sloan, & Wallis, 2013; Kennerley, Dahmubed, Lara, & Wallis, 2009; Rushworth & Behrens, 2008; Walton, Kennerley, Bannerman, Phillips, & Rushworth, 2006; Walton et al., 2009; Walton, Bannerman, Alterescu, & Rushworth, 2003;. Walton, Bannerman, & Rushworth, 2002; Walton, Rudebeck, Bannerman, & Rushworth, 2007). In the last few years a similar network has been characterized in humans. Lesions of MPFC may result in a condition known as akinetic mutism (Devinsky, Morrell, & Vogt, 1995), whereby patients show difficulties in initiating speech and movement, not due to an impairment of related systems, but rather to the inability or “lack of will” to start it. Electrical stimulation of the same region seems to induce a general feeling of being more motivated and more willing to persevere in effortful endeavors (Parvizi, Rangarajan, Shirer, Desai, & Greicius, 2013, although this single-case study presents some methodological caveats). More recently, neuroimaging studies have shown involvement of the striatum and MPFC in effort-reward trade-off computations (Botvinick, Huffstetler, & McGuire, 2009; Croxson, Walton, O’Reilly, Behrens, & Rushworth, 2009; Engstrom, Landtblom, & Karlsson, 2013; Massar, Libedinsky, Weiyan, Huettel, & Chee, 2015; Mulert et al., 2008). Furthermore, expecting to perform a more cognitively challenging task is associated with increased activity in striatum and MPFC, overlapping with activity induced by the prospect of a higher reward (Krebs, Boehler, Roberts, Song, & Woldorff, 2012; Vassena, Silvetti, et al., 2014).

These results suggest a crucial contribution of MPFC to effort-based behavior, hypothesized to compute the willingness to engage in the task at hand, given that upon completion a reward will be received. This principle has been defined in recent theories (Holroyd & Yeung, 2012; Shenhav, Botvinick, & Cohen, 2013), and formalized in a computational model wherein MPFC calculates the value of boosting certain actions over others, accordingly guiding behavior in cognitive and physical tasks (Verguts, Vassena, & Silvetti, 2015). Such computations are thought to influence decision-making (Treadway, Buckholtz, et al., 2012), resource allocation, task preparation (Kurniawan, Guitart-Masip, Dayan, & Dolan, 2013; Vassena, Silvetti, et al., 2014), and response vigor (Kurniawan, Guitart-Masip, & Dolan, 2011), even at the lowest layers of the motor system (Vassena, Cobbaert, Andres, Fias, & Verguts, 2015).

In summary, growing evidence supports a pivotal role of MPFC in effort-based behavior. However, such empirical effects and theorizing efforts have so far failed to provide a precise computational characterization able to account for this line of evidence within other existing computational frameworks of MPFC function.

### The Predicted Response Outcome (PRO) Model

According to the PRO model, MPFC implements two core mechanisms: 1) learning to predict the outcome of responses generated in response to environmental stimuli and 2) signaling discrepancies between predictions and observations. Using these two primary signals as an index of MPFC activity, the PRO model has previously been shown to account for a range of effects observed in MPFC related to cognitive control and decision making, including effects of error, conflict, and error likelihood. Critically, the PRO model explains these effects without reference to the underlying affective import: feedback related to behavioral error is equivalent to feedback indicating correct behavior in the sense that both forms of feedback constitute an outcome that can be predicted on the basis of task-related stimuli. An open question, therefore, is how the PRO model might be extended to account for effects in which behavior is influenced not only by the likelihood of an event occurring, but also by the value of that event.

**Figure 1.**
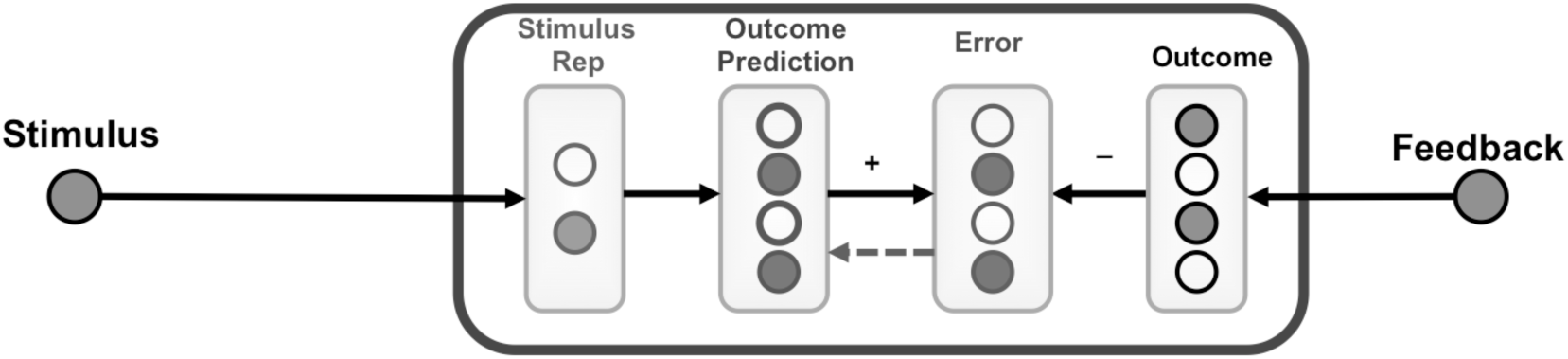
PRO model architecture (adapted from Alexander and Brown, 2011). The circles outside the box represent environmental input (stimulus and feedback). The circles inside the box represent units coding neural activity. Stimulus representations code environmental stimuli. Depending on previous occurrence, certain stimuli predict certain outcomes, as coded in outcome prediction units. Outcome units code real environmental outcome (feedback). A comparison between outcome prediction units and outcome units results in an error signal (discrepancy between predicted and actual outcome). This error signal feeds back into the outcome prediction unit, to update such predictions.

### Translating the PRO model to effort-based motivation

According to the PRO framework, MPFC activity encodes prediction error, resulting in increased activity for more unexpected (surprising) events. However, several studies investigating effort-based behavior report increased activity in the same region of MPFC when more effort needs to be invested (i.e. in presence of a more demanding task, Krebs et al., 2012; Vassena, Silvetti, et al., 2014). To reconcile this apparent inconsistency, we hypothesize that MPFC contribution to effort-based decision making parallels its role in cognitive control – MPFC predicts the amount of effort (as well as reward) associated with certain environmental cues, and the likelihood of the choice to engage or not in the required behavior. In other words, we propose that MPFC monitors effort cues and decisions, with the same mechanisms used to monitor the occurrence of any other stimulus and response outcome.

Decisions regarding whether to engage in an effortful task carry multiple consequences. First, the choice to perform an effortful task entails exerting actual effort in order to perform the task (regardless whether the task is performed successfully or not). Additionally, performing a task carries with it the possibility of success, in which case the subject receives positive feedback, often in the form of monetary reward. Alternately, the subject may fail to perform the task successfully, in which case negative feedback is provided indicating failure. In the simulations, such failure corresponds to not realizing the monetary reward, rather than losing money (although a loss condition could also be simulated as easily). In the framework of the PRO model, then, the outcomes predicted during choices regarding whether to engage in an effortful task are 1) the level of effort to be exerted and 2) the potential expected payoff. Furthermore, our implementation relies on two assumptions. First, greater effort is considered an aversive outcome, which generally tends to be avoided if possible (Kool et al., 2010). Second, as in the original model implementation (Alexander & Brown, 2011), outcomes can be more or less salient: increasing levels of reward and effort correspond to increasing salience in the model. This assumption is based on the observation that effort is frequently perceived as aversive, plausibly generating increased arousal level.

Under these assumptions, we simulated effort-based decision making with the PRO model. The parameter set used here was the same used in simulations reported in earlier work (Alexander & Brown, 2011, 2014), with no additions to the architecture of the model, and therefore not specifically tailored to the current context (the code is available at https://github.com/modelbrains/PRO_Effort). One should note that in this case the PRO model is not performing the task itself, but rather monitoring the choice of engaging in more or less effortful and rewarding trials (i.e. updating its predictions as a function of the experienced effort and reward, as if the task had been performed), as opposed to accepting a default option with a low reward value and no effort. In this formulation, MPFC activity reflects a monitoring signal, tracking the (un)predictability of motivationally relevant variables, instead of explicitly computing a cost-benefit trade off or driving choice. Related work (cf. Brown & Alexander, this issue) suggests how signals generated by the PRO model may be deployed elsewhere in the brain to drive choice behavior. Additionally, the adaptation of the PRO model to the context of effort-based decision-making suggests that the role of MPFC is primarily in monitoring the level of prospective reward and effort, and does not necessarily drive decisions to engage in a proposed task, nor, once engaged, to maintain performance levels sufficient to realize successful completion of a task. Rather, according with additional applications of the PRO model in this issue (cfr. Brown & Alexander, this issue), signals generated by MPFC are incorporated into decision processes occurring beyond cingulate itself. This interpretation of MPFC function is at odds with other models of MPFC function (Holroyd & Yeung, 2012; Shenhav et al., 2013), and provides a novel view of the role of MPFC in motivated behavior that may be the target of future research.

For simulations of the effort-based decision-making task, the model was presented a compound cue indicating the level of prospective reward (4 levels) and level of prospective effort (4 levels). Each reward level was modeled as a single input unit, as was each effort level, for a total of 16 unique compound stimuli reflecting combinations of effort and reward information. Following a decision to perform the task, the model received feedback related to the level of reward received and the level of effort expended. The strength of the feedback signal for both effort and reward was set to the level indicated by the corresponding model input (1 through 4) multiplied by a constant (0.48 for reward, 0.55 for effort). The constant was selected by hand to reproduce the qualitative pattern of behavioral effects reported in the literature (Klein-Flügge, Kennerley, Saraiva, Penny, & Bestmann, 2015). Following a decision not to engage in the effort task, feedback was set to 1/4 of the lowest reward level.

Figure 2 shows the results of the simulations. The model behavior replicates qualitatively effort avoidance tendencies of human participants (see Figure 2a): as the required effort (task difficulty) increases, the probability of engaging in the task decreases (i.e. the prediction that one will choose to engage). Plausibly, the prospect of a high reward changes this pattern: when a higher reward is expected, the probability of engaging in more effortful tasks decreases only slightly relative to low reward conditions. These behavioral predictions are consistent with empirical findings of previous studies (Apps et al., 2015; Klein-Flügge, Kennerley, Friston, & Bestmann, 2016). By looking at activity of the prediction units in the PRO model (Figure 2b), one can extrapolate quantitative predictions about expected activity in MPFC across different effort and reward conditions. According to the simulation, MPFC activity monotonically and linearly increases as a function of increased required effort (task difficulty) when reward prospect is high. However, when reward prospect is low, MPFC activity increases less steeply and only up to a certain degree of required effort, subsequently decreasing as the probability of engaging in trials with high-demand for low reward drops. To our knowledge, this neural prediction is yet to be tested and could be investigated by recording MPFC activity during effort-based decision-making when difficulty is manipulated parametrically.

**Figure 2.**
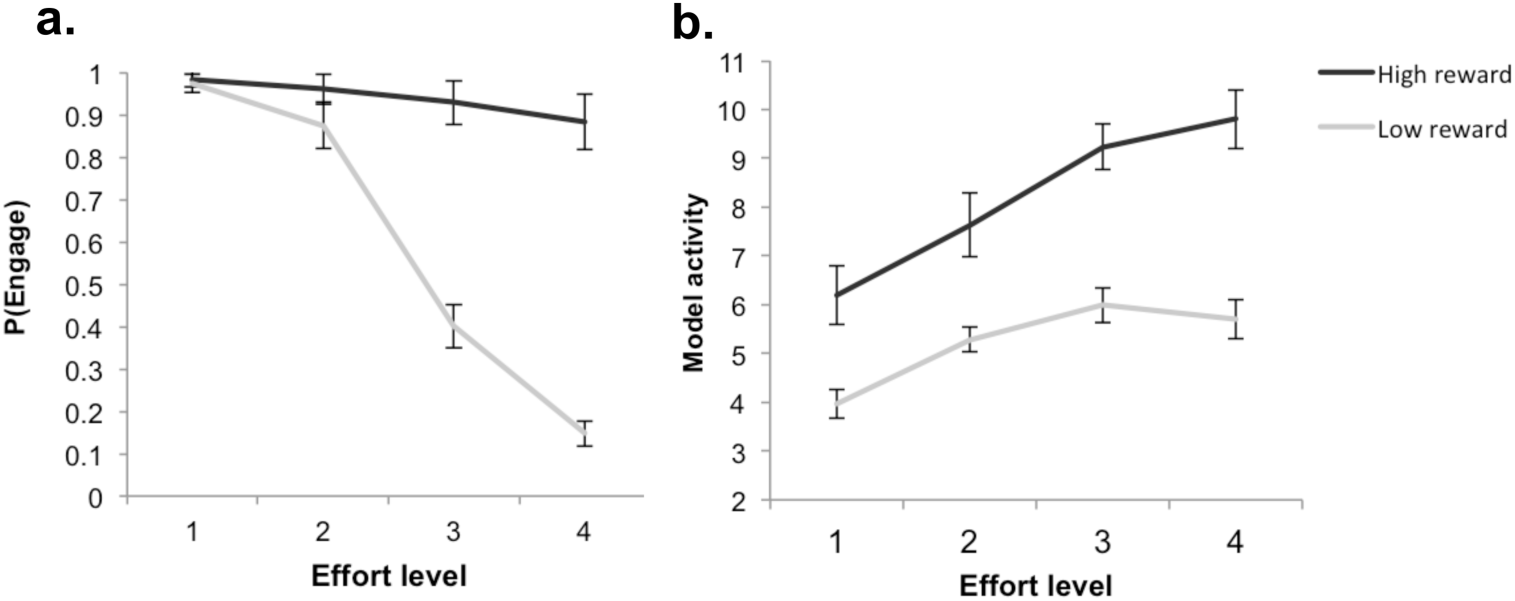
Model predictions. a. Behavioral predictions. The y-axis shows the probability of choosing to engage in a task. The x-axis shows four different effort levels (varying parametrically from easy (level 1) to hard (level 4). The grey line indicates a low reward upon successful completion. The black line indicates a high reward upon successful completion. The plot shows that with a low reward, increasing task difficulty reduces the probability of engaging in the task, while with a high reward the model engages in the increasingly effortful task anyway. b. Neural predictions. The y-axis shows MPFC activity at the time of cue. The x-axis shows the four effort levels. The grey line indicates low reward. The black line indicates high reward. The plot shows that model activity is overall higher when reward is high. Moreover, when reward is high activity linearly increases as a function of increasing effort. When reward is low, model activity only increases up to effort level 3.

### Alternative models of effort-based behavior

Other theoretical and computational models have been developed to account for MPFC contribution to effort-based behavior (Shenhav et al., 2013; Verguts et al., 2015). These models present one major difference with respect to the PRO framework: they explicitly operationalize effort as a cost to be computed in MPFC. As a result, while these models work well in predicting effort-based decisions and task-performance, they do not provide an explicit computational characterization of how MPFC contributes to other empirical effects.

Verguts and colleagues (2015) assign MPFC a role in calculating the benefit of deploying effort in addition to signaling potential rewarding outcomes. Their adaptive effort investment model operationalizes effort explicitly by implementing what the authors call “boosting”. In this model, units representing MPFC activity compute the value of boosting, namely exerting the effort needed to energize a more difficult action (be it a physical action or a cognitive task). Boosting, as in exerting effort, entails a cost. If the value of boosting outweighs the cost, the more effortful action will be selected. This results in the following predicted pattern of activity: overall activity in MPFC should be higher for larger rewards, increase with increasing task-difficulty as long as the reward is worth the effort, and drop for tasks too difficult to be solved. To our knowledge, this prediction still requires empirical testing. In line with this model, Shenhav and colleagues proposed the “expected value of control theory” (EVC, Shenhav et al. 2013).This theoretical framework assigns MPFC the role of computing the value of exerting control, by combining “component computations” estimating costs, benefits, and consequences associated with control signals (Shenhav, Cohen, & Botvinick, 2016). Input signals to such computations may include error, conflict, difficulty and prediction errors signals, which may originate outside MPFC.

From the implementation point of view one should consider that the adaptive effort allocation and EVC frameworks rely on very different assumptions as compared to the PRO model. The first two place computation of the value of boosting (for cognitive or physical action in Verguts et al.) or exerting control (cognitive tasks in Shenhav et al.) in MPFC, while the PRO model places prediction and prediction error computations in MPFC. Moreover, whereas the PRO model postulates a shared underlying computational principle, adaptive effort allocation and EVC imply the coexistence of different computations (cost and value of boosting, and prediction error for the first, separate cost, benefit and consequences estimation for the second). However, further modeling work is required to extrapolate predictions, which may disentangle the models based on available empirical evidence.

The main advantage of the PRO model is parsimony: the same architecture explains effort-related effects as well as a wide variety of empirical effects previously measured in MPFC (ranging from prediction error, cognitive control, conflict and so forth, Alexander & Brown, 2011). This is not the case for the adaptive effort investment model, which is specifically tailored for effort-based behavior and is therefore not applicable in other contexts, at least in its current implementation.

One limitation of the PRO model is that it does not perform the task and is not responsible for action selection: MPFC units compute predictions and compare them with outcomes. This assumes that the reward and cost trade-off computations, and the choice to engage or not in the task at hand are implemented elsewhere. Candidate areas could potentially be other sub-regions of PFC, or possibly the basal ganglia and especially the striatum, shown in several studies in both humans and animals to contribute to effort-based decisions and task-preparation (Bailey, Simpson, & Balsam, 2016; Botvinick et al., 2009; Prévost, Pessiglione, Météreau, Cléry-Melin, & Dreher, 2010; Salamone, Correa, Farrar, & Mingote, 2007; Vassena et al., 2014). One shortcoming common to both PRO and EVC/adaptive effort allocation frameworks is that they are agnostic about cost computation. Effort is plausibly defined as a function of task-difficulty and higher effort equals higher cost. However the nature and source of such cost signal, is a topic of ongoing empirical and theoretical work (Holroyd, 2016; Kurzban et al., 2013).

### Predictions and implications for clinical populations

Adaptive decision-making and energization of behavior poses a challenge in several daily life situations. In a number of psychiatric conditions, these mechanisms are impaired. Recent studies showed that decision-making regarding whether to undertake an effortful task is altered in depression, bipolar disorder and schizophrenia (Barch, Treadway, & Schoen, 2014; Culbreth, Westbrook, & Barch, 2016; Hershenberg et al., 2016; McCarthy, Treadway, Bennett, & Blanchard, 2016; Silvia et al., 2016; Silvia, Nusbaum, Eddington, Beaty, & Kwapil, 2014; Treadway, 2016; Treadway, Bossaller, et al., 2012; Yang et al., 2014). Symptoms often include reduced willingness to exert effort, although data across different pathologies or effort types do not always align. For example, both schizophrenia and depressed patients show reduced allocation of physical effort for higher rewards as compared to controls, while evidence concerning cognitive effort is mixed (Barch, Pagliaccio, & Luking, 2016). The same authors suggest a different underlying deficit for the two conditions: depressed patients seem to show impaired hedonic processing, while schizophrenia patients tend to show impaired reinforcement learning and action selection. Moreover, effort-related deficits in schizophrenia point to an effort allocation deficit, rather than reduced effort expenditure per se (McCarthy et al., 2016; Treadway, Peterman, Zald, & Park, 2015), with patients performing less optimal decisions. Such a complex picture confirms alteration of effort-based decision-making in such clinical populations, and calls for more precise quantitative frameworks, able to identify the mechanisms underlying different impairments.

Here, we use the PRO model, adapted as described above for modeling effort-related dynamics in healthy subjects, to simulate the possible neuroetiology underlying clinical disorders, which could explain the behavioral symptoms measured in clinical samples. In the PRO model, outcome representation units may be modulated by salience (Alexander & Brown, 2011) suggesting that compromised function in clinical populations may be a result of altered perception of salient events (Alexander, Fukunaga, Finn, & Brown, 2015). Model simulations and theoretical predictions are described in Figure 3.

**Figure 3.**
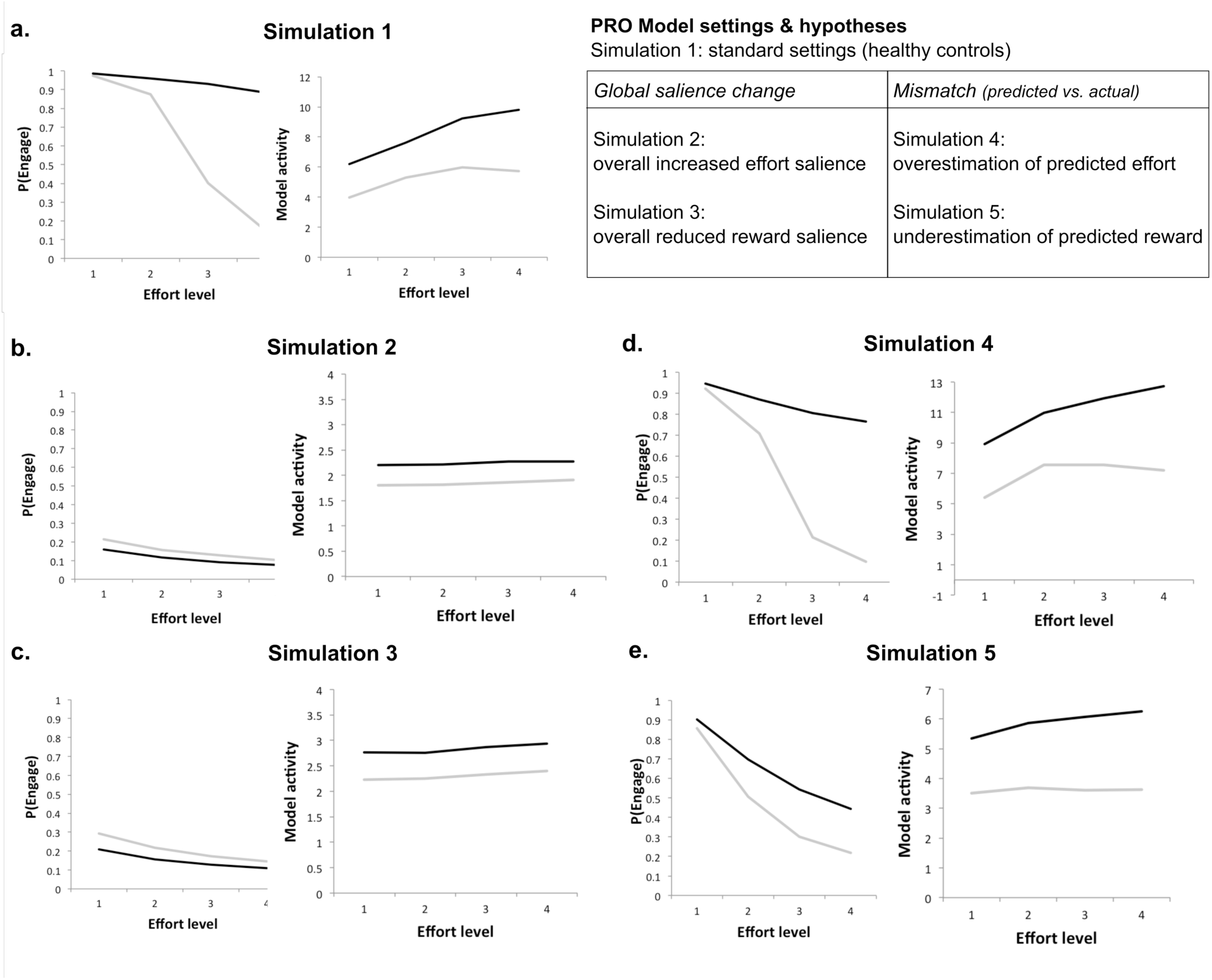
PRO model simulations of impaired motivation. In all plots, the y-axis shows the probability of engaging in the task (left panel) and the model activity (right panel). The x-axis shows four possible effort levels, parametrically increasing from easy (level 1) to hard (level 4). Grey lines indicate low reward upon task completion. Black lines indicate high reward upon task completion. a. Simulation 1. Behavioral and neural predictions for healthy controls. The table on the right illustrates the hypotheses of possible impairments as modeled with the PRO model, and relative explanation. We hypothesize two core possible mechanisms driving impairments in patients. The first is altered *global salience*, with either an overall increased effort salience (simulation 2), or an overall increased reward salience (simulation 3). The second is *mismatch* between predicted and actual outcome, with either a possible overestimation of predicted effort (as compared to actual, simulation 4) or a possible underestimation of the predicted reward (as opposed to actual, simulation 5). b-e. Simulations 2 through 5. Behavioral (choices) and neural (model activity) predictions under the different hypotheses.

These simulations use the basic architecture of the PRO model without modification as in simulation 1. In order to simulate altered function during effort-based decision-making, we assume that clinical disorders entail alterations in the processing of information related either to reward or effort information. One possible alteration driving impairment in decision-making could be attributed to a *global salience change*: in some populations, the global salience of decision variables might be affected. Patients may be overly sensitive to the costs of engaging in a task (such as required effort, simulation 2, Figure 3b), or have reduced sensitivity to potential reward (simulation 3, Figure 3c). To simulate these hypotheses, we multiply the effort level from simulation by a factor of 2 (simulation 2), or the reward information by a factor of 0.5 (simulation 3) to reflect increased effort salience or decreased reward salience. The results of these simulations show that increasing the salience of effort and reducing the salience of reward have similar effects in the model: the probability of engaging in a task is decreased over all levels of reward and effort. The pattern of MPFC activity predicted by the model is also severely attenuated relative to control simulation: activity is slightly higher in the high reward as compared to low reward condition, but does not seem to track effort as it did in the control simulation.

Another possible alteration underlying the impairment in clinical populations might be a *mismatch*: predictions made by the model regarding effort and reward levels might not correspond to veridical experience. The model may overestimate the level of effort required (simulation 4) or underestimate the value of the reward on offer (simulation 5). The inability to accurately estimate required effort and potential reward, would generate a mismatch between prediction and outcome: predicted effort could be overestimated, leading to abnormal effort avoidance, while mismatches between predicted and experienced reward could lead to decreased motivation in performing the task. To simulate these hypotheses, effort-related feedback to the model was multiplied by a factor of 2 (simulation 4), while the valence information used for updating top-down control weights (Alexander & Brown, 2011, supplementary Figure 1) remained unchanged. The net effect is that the model’s prediction of effort level exceeds the effort experienced by the model following choices to engage in an effortful task. In simulation 5, reward-related feedback to the model was multiplied by 0.5 (while valence information was unchanged), with the interpretation that the level of predicted reward did not match the experienced level. Simulation results for effort mismatch (Fig. 3d) and reward mismatch (Fig. 3e) show that such mismatches in effort and reward prediction yield qualitatively different predictions regarding behavior: overestimation of effort level leads to increased discounting of low reward offers, while behavior in high-reward conditions is relatively unaffected compared to control simulations. Conversely, underestimation of reward produces a general increase in discounting - both high and low reward conditions are discounted more heavily compared to control simulations.

To our knowledge, the hypothesized mechanisms (*global salience impairment vs. predicted/actual mismatch*) cannot be disentangled on the basis of existing data. Future experimental work to test this may incorporate a model-based fMRI experiment with patients performing an effort-based decision-making task. One could simulate model-based predictors of MPFC activity for each hypothesized mechanism on the basis of subjects’ actual performance. This would show which mechanisms better explain brain activity measured in MPFC (i.e. the one giving better fit between model activity and neural data). Empirical verification in clinical populations showing impairments of effort-based behavior would shed light on potential mechanisms underlying symptoms origin and provide (in)validation for the PRO as plausible neurofunctional account of MPFC contribution to motivated behavior.

### Limitations and critical aspects

Translating the PRO framework to a motivational context allows explaining effort-based behavior under a working computational model of MPFC functioning without postulating a MPFC function dedicated to explicit cost computations. However, this translation leaves some critical aspects unanswered, open for future research. First, in our conceptualization we do not distinguish between different types of effort costs, such as physical vs. cognitive effort. Here we only assume higher effort to be more salient and aversive, irrespective of its specific nature.

Previous research comparing neural circuits involved in different effortful tasks (Schmidt, Lebreton, Cléry-Melin, Daunizeau, & Pessiglione, 2012) suggests that the type of effort determined the network involved in task execution, with motor regions implicated in a physical task as opposed to parietal regions implicated in a cognitive task. In both cases, the relevant network was more active in the high effort condition. Moreover, a shared motivational hub was identified in the striatum, showing increased activity irrespective of effort type. In both animal and human research the striatum has been implicated in cost-benefit trade-offs (Botvinick et al., 2009; Salamone et al., 2016; Vassena, Silvetti, et al., 2014; Westbrook, A. & Braver, 2016), and is often co-active with MPFC. An intriguing possibility is that striatal dopamine-driven trade-off computations provide MPFC with the necessary cost signal regulating subsequent behavior, irrespective of effort type. These speculations should be investigated in further research.

Second, we do not include a mechanistic explanation of the aversive nature of effort. The neural origin of this computation is still debated in the literature. It has been proposed that perception of effort cost derives from its opportunity cost (i.e. engaging resources which could be utilized differently, Kurzban et al., 2013). A recent account hypothesizes effort cost to derive from accumulation of waste product at the neural level, resulting from using up neural resources (Holroyd, 2016). The model is currently agnostic to the origin of this signal, which we consider an avenue for future modeling and experimental work.

Third, we formulated effort-based behavior as a decision-making problem, where effort and reward are considered outcomes of the decision to engage in the task at hand. However, this does not account for monitoring ongoing effort exertion. Maintaining a certain level of vigor throughout a period (e.g. holding a grip) could be seen as the result of a series of decisions to keep engaging throughout the entire period, depending on (presumably striatal) cost and reward signals fed into MPFC. This intriguing idea should be addressed in future modeling and experimental work.

Fourth, we do not simulate MPFC activity variations within a trial. Theoretically, the PRO model states that MPFC continuously predicts stimulus-outcome associations (Alexander & Brown, 2014). This means that at the beginning of a trial, prior to effort or reward related information being presented, the model would predict average outcomes (in the context of effort-based decision-making, these predictions would converge on the mean reward and effort for the overall task). Following cue presentation, this prediction would be updated when experiencing the actual effort, suggesting that MPFC activity should reflect the degree by which task-related cues on a specific trial diverge from the average experimental value. Preliminary evidence for such a computation is reported in a Transcranial Magnetic Stimulation study measuring motor-evoked potentials (Vassena et al., 2015), which showed that motor cortex excitability during cue presentation was related to prediction error in expected value (discrepancy between average expected value and value of the actual cue on the current trial, integrating a certain degree of required effort and potential reward). However, how this result speaks to MPFC contribution in the process remains to be investigated. Conversely, activity following the choice regarding whether to engage with an effortful task should vary inversely with the tendency of the subject to engage: subjects with a lower overall tendency to engage in effortful tasks should show increased MPFC activity following choices to engage, while subjects with a strong tendency for engaging should show increased MPFC activity following choices not to engage. These theoretical predictions require empirical testing, and possibly additional modeling work to specify them quantitatively. From the methodological point view, one would need to collect fMRI data at a time scale with sufficient resolution to contrast MPFC activity at both cue and outcome, or to use EEG-fMRI simultaneous recordings to localize MPFC electrophysiological signature.

## Effort-based decision-making and performance in DLPFC

In the existing literature, the link between DLPFC and effort-based behavior is more implicit, although it clearly emerges from the number of high-level functions implicating this region. Traditionally, DLPFC is assigned a pivotal role in supporting working memory updating and maintenance (Curtis & D’Esposito, 2003; Miller & Cohen, 2001). In recent years, evidence has accumulated showing a crucial contribution of this region to executive functions, including goal-maintenance and task-set representation (Ridderinkhof, van den Wildenberg, Segalowitz, & Carter, 2004). Activity in DLPFC is associated with maintaining stimulus representations and strategies for optimal task performance. Although several frameworks have been proposed to explain DLPFC function (Badre, 2008; Koechlin, 2014), a mechanistic account of how such representations and strategies are formed and manipulated to guide goal-directed behavior is still lacking. How DLPFC interacts with MPFC prediction signals remains unclear. Several studies investigating motivation and task-preparation report co-activation of DLPFC and MPFC (Botvinick & Braver, 2015; Chong et al., 2017; Engström, Karlsson, Landtblom, & Craig, 2014; Engstrom et al., 2013; Krebs et al., 2012; Rypma, Berger, & D’Esposito, 2002; Vassena, Silvetti, et al., 2014). Across these studies, DLPFC activity increases as a function of expected effort, task-load and working memory demands. Recently, starting from the principles outlined in the PRO model, it has been proposed that the underlying computational mechanism of DLPFC might also rely on the prediction and prediction error (Alexander & Brown, 2015). These authors proposed an updated version of the PRO model, extended in a hierarchical architecture: the Hierarchical Error Representation model (HER).

**Figure 4.**
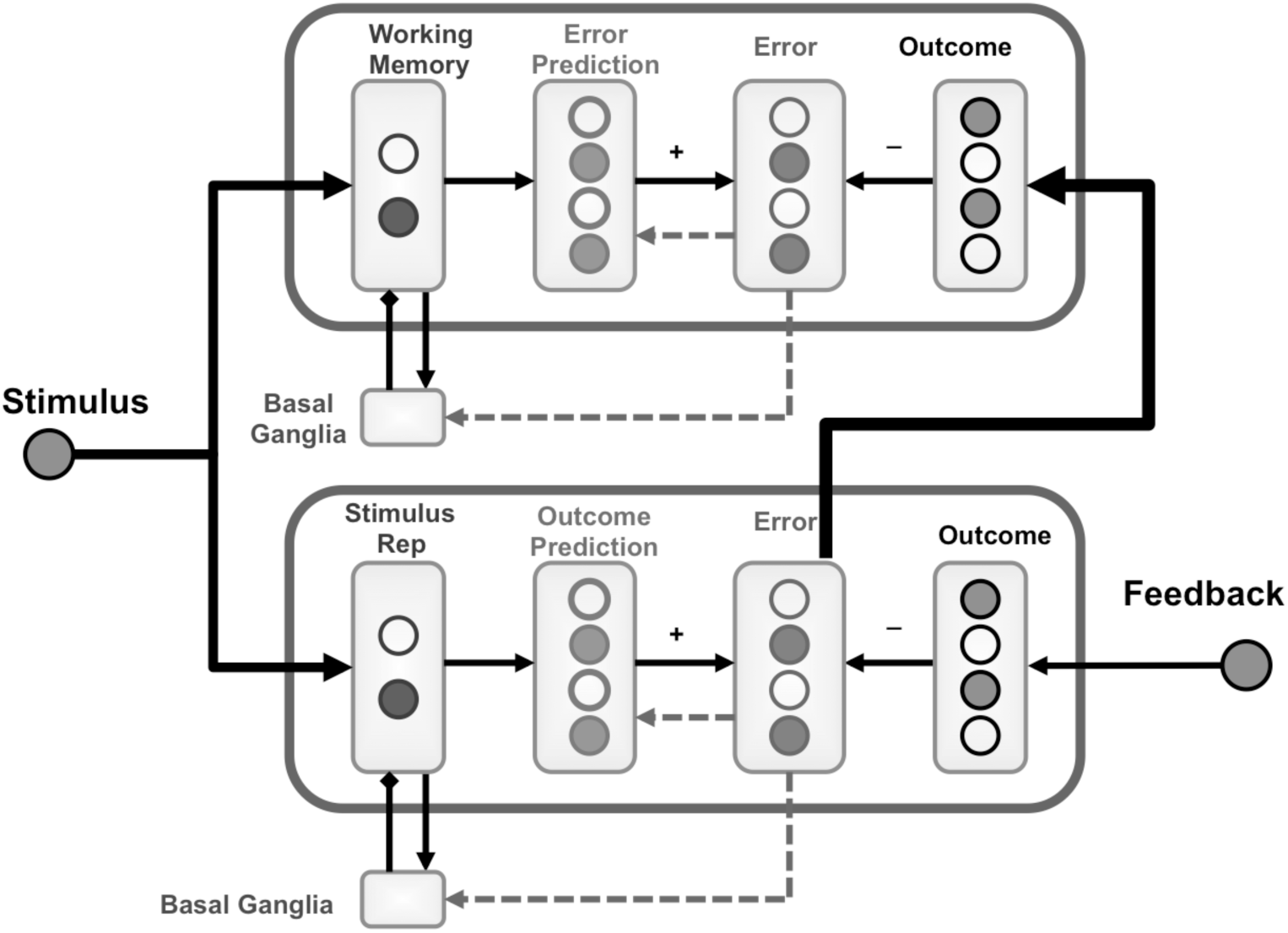
HER model architecture (adapted from Alexander and Brown, 2015). The circles outside the box represent environmental input (stimulus and feedback). The circles inside the box represent units coding neural activity. This figure shows a 2-layer version of the HER model. Each layer replicates the architecture of the PRO model (cfr. Figure 1): stimulus representation unit code for environmental stimuli, leading to prediction of a certain outcome (outcome prediction units); outcome units code for actual outcome; comparison between outcome prediction and actual outcome produces an error signal, coded in the error units. In the HER model, the activity of error units at the lowest layer scales with the discrepancy between predicted and actual outcome (as in the PRO model); error signals update predictions in this layer, but are also fed to the upper layer, where actual error signal is compared with the predicted error signal. The resulting 2^nd^ level error signal is used to update predictions of the future error signal.

The HER model is composed of 2 or more hierarchical layers, and each layer replicates the functional form of the PRO model. The lowest layer receives input and feedback from the environment, updating predictions via prediction error, computed as the discrepancy between predicted and actual outcome. Additionally, the error signal also provides input to the layer above, where it is treated as a feedback signal; in other words, this higher layer learns predictions of the expected error of the lower layer, compares such prediction with the actual error signal, and updates the error prediction accordingly. This simple architecture provides a mechanistic account of how MPFC and DLPFC might interact, congruent with available empirical evidence (Alexander & Brown, 2015, 2016). The prediction error signal generated in MPFC not only results in an updated error prediction at the highest layer: this prediction is also linked to the environmental stimulus (or context), which was associated with the error. This results in a representation linking the error signal to the stimulus (or context) that preceded the error. In agreement with a substantial body of evidence, this model accounts for the primary role of MPFC in performance monitoring and error detection, and for the role of DLPFC in maintaining task-set representations providing context for MPFC function.

### Future directions: translating the HER model to effort-based behavior

Despite its wide explanatory power, the HER framework has to date not been translated to the domain of motivation to accommodate for the aforementioned effort and task-load effects observed in DLPFC. In the previous sections we showed the potential of the PRO model to explain effort-related effects. Fundamentally, the HER model is an extension of the PRO model, which suggests it might be well suited for a comparable translation to the effort domain.

The aim of the current section is twofold. First we propose a theoretical explanation of how DLPFC-MPFC interaction in the context of the HER model could account for motivational effects observed in both regions. Second, we derive informal behavioral predictions from the HER model in its current formulation which can be tested in both healthy and clinical populations to further challenge the validity of the model. One should note that such interpretations and prediction are highly speculative at this stage. The purpose of this section is to provide a series of directions and predictions to drive empirical investigation of DLPFC-MPFC contribution to effort-based behavior.

The HER model is built on the principle that error signals in MPFC are equivalent to other environmental feedback signals, and are therefore subject to the same prediction and error processes. When an error signal is unexpected, the error prediction is updated. This *error history* is stored in DLPFC as error representations linked to stimuli or environmental contexts. This implies that when the same stimulus or environmental context reoccurs, the corresponding DLPFC error representation is also reactivated. We hypothesize that this representation will in turn up-regulate MPFC activity, reinstating the signal experienced at the time of error, but this time with the purpose of exerting control to prevent the error from happening again (thus leading to a better prediction, or a successful behavioral outcome). In this formulation, the translation to a motivational context becomes evident: a performance error, for example due to task difficulty, would be signaled by increased MPFC activity, tagging that particular behavioral instance as requiring extra effort. Next time the same instance reoccurs, the reactivated error representation can provide information necessary to inform top-down control and resource allocation to result in successful task performance. Noteworthy, this speculative explanation does not require an explicit operationalization of effort or other motivational factors: thus, the HER model in its current architecture could be able to account for both prediction-related as well as effort-related signals in MPFC and DLPFC. The empirical validity of this explanatory framework is to be tested in future research, which should provide neurobiological evidence for the type of MPFC-DLPFC dynamics described above.

Besides the theoretical implications for understanding PFC circuitry, the model relies on assumptions that require empirical testing. The hierarchical structure of the HER model is consistent with other accounts of PFC function, postulating the existence of a cortical rostro-caudal hierarchical gradient (Badre, 2008; Koechlin, 2016). According to these theories, caudal regions of PFC encode more concrete representations (action-related, or more recent in time), while more rostral regions encode more abstract representations (task-sets, rules, context or information further in the past to be maintained). This is implemented in the HER model, wherein a typical simulation of a working memory task, different items to be stored in working memory are encoded at different levels of the hierarchy (depending on order of processing, see for example the 12AX task simulations in Alexander and Brown, 2015). An underlying assumption is serial processing, not only for series of stimuli, but also for complex stimuli composed by different stimulus features. When placed in the context of motivation and decision-making literature, this assumption is quite relevant: most of the experiments referenced above combine motivationally salient information of different types, such as required effort and available rewards, presenting it simultaneously. The empirical question remains open as to whether such simultaneous presentation results in simultaneous or serial processing of the presented information sources, and to date this question has not been addressed. The HER framework hypothesizes that such features would be processed serially in a specific and preferred order. Simulations showed that altering this order, by imposing a non-preferred order, can impact performance (Alexander & Brown, 2015).

Presenting motivationally salient information prior to task performance typically influences accuracy, reaction time and task preparation in several tasks requiring cognitive control (Aarts & Roelofs, 2010; Boehler, Schevernels, Hopf, Stoppel, & Krebs, 2014; Janssens, De Loof, Pourtois, & Verguts, 2016; van den Berg, Krebs, Lorist, & Woldorff, 2014; Vassena, Silvetti, et al., 2014). When applied to the domain of effort-based behavior, the order hypothesis predicts that altering order of processing of reward and effort information might result in a shift in perceived subjective value, and consequently affect (improve or deteriorate) performance.

### Predictions and implications for clinical populations

These theoretical predictions naturally stem from the HER model, and empirical testing of their validity carries relevant implications. First, testing these predictions will (dis-)prove the validity of the assumptions underlying the model. Second, if altering order of processing can alter decision-making, one could test the potential of such manipulation to improve dysfunctional decision making, for example concerning health-related behavior such as physical exercise and eating habits. Third, if altering order of processing can alter performance, one could devise optimal ways to reconfigure available motivational information to improve cognitive performance, for example in educational and school settings. Lastly, all of the above have important implications for translational research and potential applications in clinical populations affected by disorders of motivation.

To date, the predictions listed above have not been empirically tested. It is however useful to speculate on the mechanisms, which could underlie such effects. One plausible explanation involves salience. If effort and reward information is processed serially, the order of processing when presentation is simultaneous may be influenced by the respective salience of informative cues. Patients with depression typically show reduced willingness to exert effort to obtain a reward: in other words they are more effort-avoidant as compared to controls (Treadway, Buckholtz, Schwartzman, Lambert, & Zald, 2009; Yang et al., 2014). One possible reason could be the overestimation of the amount of the required effort, which would result in an unfavorable overall value, leading to the decision of not engaging in the task. Similarly, reward information could be underestimated, thus reducing the worth of the final value. Note that these hypotheses are in line with what was formulated and simulated with the PRO model in the previous section of this manuscript, where we hypothesized impairment in perceived salience of effort and reward. What is particularly relevant with respect to the order effects, is the possibility of intervention: motivational impairment could derive from altered perception of saliency; manipulating order of presentation may enforce a specific order of processing, artificially increasing or decreasing saliency of effort and/or reward information; by tuning this manipulation, one might be able to determine the optimal configuration restoring normal perception of salience. Ideally, this process would result in increasing the willingness to exert effort in exchange for reward, thus counteracting the typical behavioral pattern of anhedonia, a core symptom in depression (Silvia et al., 2014; Treadway, Bossaller, et al., 2012). Critically, alterations in effort- and reward-based decision making have also been reported in other psychiatric conditions such as bipolar disorder and schizophrenia (Barch et al., 2014; Fervaha et al., 2013; Gold, Waltz, & Frank, 2015; Hershenberg et al., 2016) and pre-clinical traits of apathy (Bonnelle et al., 2015), although showing different patterns of impairment. Testing the predictions derived from the HER model across different clinical samples could provide insights on shared and dissociable underlying etiopathogenetic mechanisms; moreover, such deeper theoretical understanding could foster development of behavioral treatments aimed at improving decision-making and behavioral outcomes for these patients in daily life.

## General discussion

This manuscript reviews the theoretical frameworks provided by the PRO and HER models, modeling the neurofunctional architecture of MPFC and DLPFC. Such models have originally been developed based on the core principles of prediction and prediction error to explain empirical effects found in these regions. Here we discussed how these models may generalize to the domain of motivation, focusing on effort-based behavior. We show that effort effects in MPFC can be successfully accounted for by the PRO model, which provides further predictions regarding behavior and neural activity in both healthy and clinical populations. Furthermore, we discuss the potential translation of the HER model to the domain of effort-based behavior, which accounts for empirical effects measured in DLPFC, and provides interesting empirical predictions regarding the effect of order of processing on decision-making and task-performance: if these predictions are borne out, such effects could lead to the development of useful interventions to influence altered perception of salience of effort and reward information in clinical population, potentially improving abnormal behavior.

One primary goal of this manuscript is to emphasize the importance of exploiting precise theoretical frameworks to derive predictions to test experimentally. The first advantage of such mathematically precise frameworks resides in the ability to explain several behavioral and neural effects observed in a brain region under the same computational principle. The second advantage is the possibility to generate new predictions based on the same model, which can translate to contexts to date untested or different populations. This feature is particularly useful to guide further theory-driven empirical inquiry. In a scientific age where empirical tools proliferate, basing experimental research on strong *a priori* hypotheses has become a necessary condition to allow drawing statistically meaningful and generalizable conclusions. Finally, such theoretical rigor and quantitative predictive precision provide a great tool to test potential translational applications, with broad explanatory power for understanding the neurobiology of disease.

## Acknowledgements

WHA and JD were supported by FWO-Flanders Odysseus II Award #G.OC44.13N. EV was supported by the Marie Sklodowska-Curie action with a standard IF-EF fellowship, within the H2020 framework (H2020-MSCA-IF2015, Grant number 705630).

